# Dynamics of Mitochondrial NAD^+^ Import Reveal Preference for Oxidized Ligand and Substrate Led Transport

**DOI:** 10.1101/2022.11.04.515224

**Authors:** Shivansh Goyal, Xiaolu A. Cambronne

## Abstract

SLC25A51 is a member of the mitochondrial carrier family (MCF) but lacks key residues that have been attributed to the mechanism of other nucleotide MCF transporters. Thus, how SLC25A51 transports NAD^+^ across the inner mitochondrial membrane remains unclear. To elucidate its mechanism, we used Molecular Dynamic simulations to study reconstituted SLC25A51 homology models in lipid bilayers. We observed spontaneous binding of cardiolipin phospholipids to three distinct sites on the exterior of SLC25A51’s central pore and found that mutation of these sites impaired transporter activity. We also observed that stable formation of the required matrix gate was controlled by a single salt bridge. Using simulation data and in-cell activity assays we identified binding sites in SLC25A51 for NAD^+^ and showed that its binding was guided by an electrostatic interaction between NAD^+^ and a negatively charged patch in the pore. In turn, interaction of NAD^+^ with interior residue E132 guided the ligand to dynamically engage and weaken the salt bridge gate, representing a ligand-induced initiation of transport.

**Significance:** NAD^+^ is an intermediary metabolite whose multiple functions are entwined with respiration, catabolism, and stress responses in cells. Previous sensor measurements had indicated that its continuous biosynthesis was required to sustain mitochondrial matrix levels in respiring cells, and SLC25A51 was identified as the required importer of NAD^+^ across the inner mitochondrial membrane. However, SLC25A51 has little homology to other nucleotide carriers at its substrate binding site. By combining modeling approaches and experimental assays, this work provides mechanistic insight into how human SLC25A51 recognizes its ligand, how the transporter can be regulated by its lipid environment, and an observation of ligand-induced gate opening. This represents the first description of the ligand binding site for an NAD^+^ mitochondrial carrier.

## Introduction

SLC25A51 is an essential gene that is ubiquitously expressed across all mammalian cells. Its protein controls concentrations of NAD^+^ in the mitochondrial matrix by directly importing the oxidized dinucleotide, and thus mammalian mitochondria depend on SLC25A51 for sustaining NAD^+^ levels (1–3). Loss of SLC25A51 resulted in loss of cellular respiration and oxidative flux of the tricarboxylic acid cycle. SLC25A51 further impacted the concentrations of NAD-derived metabolites and cofactors, including NADH and NADPH (1–3).

There is little understood, nevertheless, about the underlying mechanisms of SLC25A51 or how the transporter interacts with oxidized NAD^+^, compared to NADH or related molecules such as nicotinamide or the mononucleotide. SLC25A51 is a member of the mitochondrial carrier family (MCF) that localizes to the inner mitochondrial membrane (1–4). By homology, its structure harbors a pseudo-tri-symmetrical pore that is characteristic of the family (4–7). The trice repeated domain comprises two transmembrane helices joined by a short matrix helix parallel to the matrix membrane. Based on cytoplasmic and matrix-open structures of the related ADP/ATP nucleotide carriers, it is surmised that members of this structurally related family function through an alternating-access mechanism (5–11).

SLC25A51 has low conservation (<20%) in its amino acid sequence compared to its functional homologues in *S. cerevisiae* and *A. thaliana* that selectively import mitochondrial NAD^+^ (*SI Appendix*, Fig. S1) (4, 12, 13). In phylogenetic analyses, SLC25A51 did not cluster with other mammalian nucleotide carriers in the family (7, 14). Thus, it remains unanswered how SLC25A51 engages and transports its ligand.

In this work we used Molecular Dynamics (MD) simulations to study how SLC25A51 homology models engage the NAD^+^ ligand under the constraints of thermodynamics and in the context of a lipid environment. We tested our observations in intact cells with mutational analysis and measurements of in-situ free mitochondrial NAD^+^ concentrations using a LigA-based NAD^+^ biosensor (15). This study provides the first insight into the binding site of an NAD^+^ mitochondrial transporter, identifies a mechanism of ligand-induced transport, and reveals cardiolipin-binding as a regulator of SLC25A51 activity.

## Results

### Stable modeling of SLC25A51 in a lipid bilayer

To study how SLC25A51 recognizes NAD^+^ for import, we first generated apo models of human SLC25A51 in its outward-facing, cytoplasmic (c-state) conformation using Swiss-Model and AlphaFold2 modeling (16, 17). Both model structures adopted the characteristic MCF fold and were superimposable upon the solved bovine ANT crystal structure (PDB ID: 1OKC) (5). Using MolProbity, we determined that the quality of the equilibrated AlphaFold2 and Swiss-Model structures were in the 100^th^ and 97^th^ percentile, respectively (18). Model quality was further estimated using pLDDT for AlphaFold2 (*SI Appendix*, Fig. S2*A*) and QMEANDisCo analysis for both AlphaFold2 and Swiss-Model with global scores of 0.55 ± 0.05 and 0.56 ± 0.05, respectively (*SI Appendix*, Fig. S2*B* and *C*) (16, 19). A direct comparison, nevertheless, revealed a difference of 3.5 Å root-mean-square deviation (RMSD) in atomic positions between the structures. Closer inspection revealed several residues in the central pore that were hydrophobically buried in one model but exposed to the hydrophilic pore in the other (*SI Appendix*, Fig. S2*D*). Therefore, we decided to test both models in this study.

Each of the apo models was individually embedded in a 86 × 86 Å^2^ swatch of lipid bilayer using Charmm-GUI that recapitulated the composition of phospholipid components in mammalian inner mitochondrial membrane (POPC:POPE:CL=2:3:2) (20–24). The system was hydrated and modeled with an ionic concentration of 150 mM KCl. Each model was analyzed in triplicate, and each replicate was ~1 μs in duration (*SI Appendix*, Table S1). We observed that the mean RMSD over time among the replicates for both stabilized c-state models were 4.1 ± 0.3 Å and 2.9 ± 0.2 Å for the Swiss-Model and AlphaFold2 models, respectively (*SI Appendix*, Fig. S3*A*, *B* and Table S1). In parallel, as a control for the approach, we generated a homology model for c-state human SLC25A4 (ANT1, ADP/ATP carrier) based on the solved bovine structure (PDB ID: 1OKC) (5). The model for SLC25A4 (RMSD 2.8 Å) displayed a similar stability compared to the SLC25A51 c-state models over the course of the simulations (*SI Appendix*, Fig. S3*C*). We further calculated root-mean-square fluctuation (RMSF) values for all c-state models, and as expected, observed less flexibility in transmembrane regions compared to loop regions (*SI Appendix*, Fig. S3 *D*-*F*). Together, the data indicated that the experimental model systems were stable.

### Cardiolipin was required for SLC25A51 activity

With both the AlphaFold2 and Swiss-Model models, we observed spontaneous recruitment of cardiolipin molecules to three sites on SLC25A51 (Fig. 1*A* and Movie S1). Notably, all lipids were initially positioned at random and cardiolipin molecules were not docked to SLC25A51. We observed cardiolipin molecules arriving from different initial positions, displacing other types of phospholipids initially bound, and consistently limiting their interactions to the same three distinct sites on SLC25A51. Together, the data suggested site-specific recruitment of cardiolipin to SLC25A51.

**Fig. 1.**
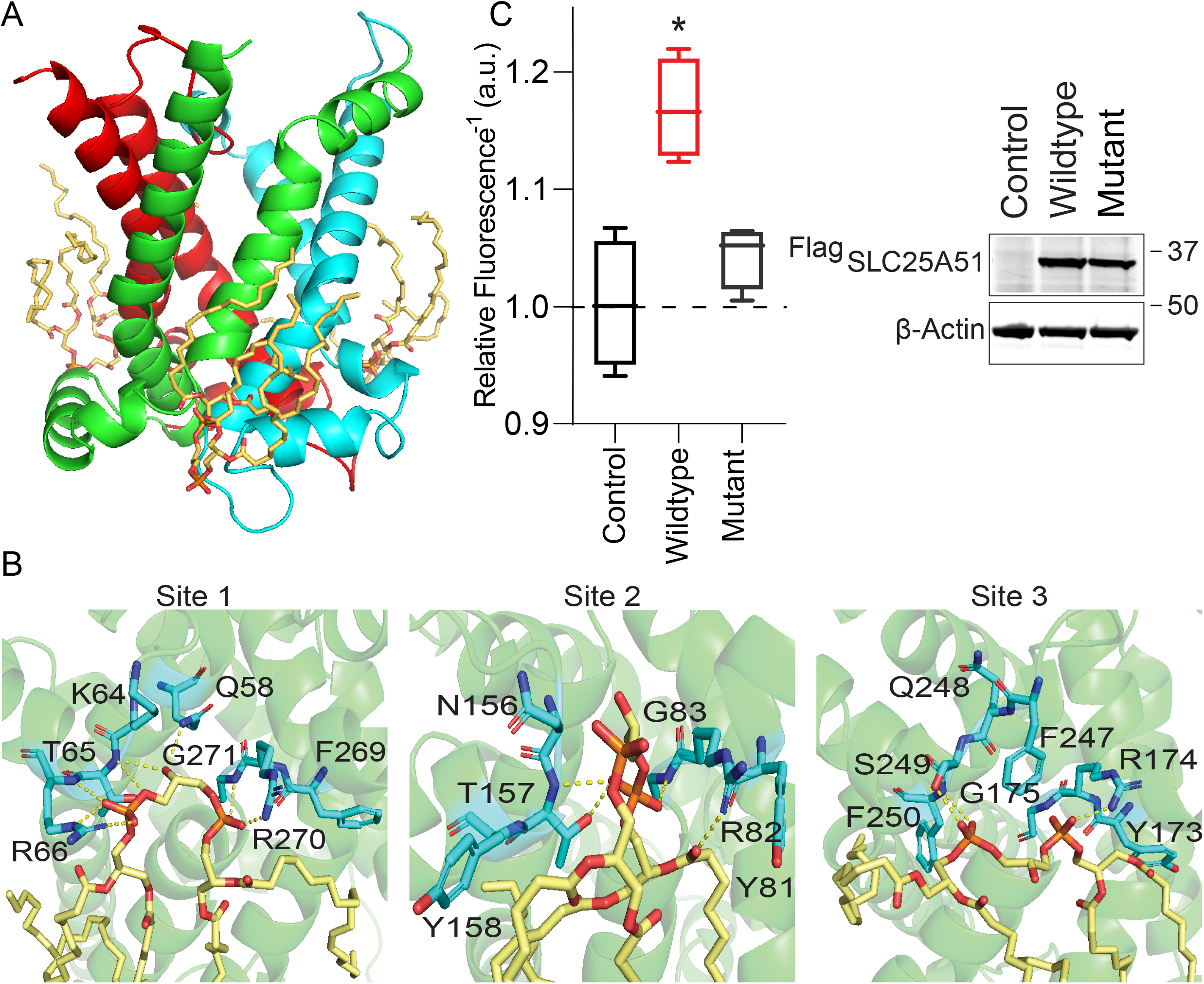
Cardiolipin binding impacts SLC25A51 function. *(A)* Observed positioning of bound cardiolipin (wheat). Differential hues in SLC25A51 delineate the three repeated domains that comprise its structure. *(B)* Representative poses depicting identified interactions at sites 1-3. Yellow dashes indicate interactions < 3.5 Å. At site 2, the binding of a single cardiolipin phosphate was often observed. *(C) (left)* Relative concentrations of free mitochondrial NAD^+^ measured using a ratiometric biosensor and ectopic expression of either Control empty vector, Wildtype ^Flag^SLC25A51, or mutant ^Flag^SLC25A51 harboring R82Q, R174Q, R270Q point mutations. Dashed horizontal line indicates baseline for unchanged ratiometric measurements at 1; mean ± SD, n = 4, ANOVA < 0.01 with post-hoc Dunnett’s Test *\*p < 0.001*. *(right)* Anti-Flag Western Blot of HeLa cells expressing indicated SLC25A51 variants or empty vector; β-actin, loading control.

Cardiolipin binding was mediated by polar interactions between its phosphates and exterior-facing residues on the transporter at Binding Site 1 (T65, R66, R270 and G271), Binding Site 2 (N156, T157, R82 and G83), and Binding Site 3 (S249, R174, G175) (Fig. 1*B*). The observed sites correlated in position with cardiolipin binding motifs [F/Y/W δ G] and [F/Y/W K/R G] identified from ADP/ATP transporter studies (5, 8, 9, 25–28). All binding sites were on the exterior of the central pore facing the matrix leaflet and positioned at the junctions between the three repeated protein domains that comprised SLC25A51’s pore (Fig. 1*A* and *B*). Each cardiolipin molecule spanned an even-numbered helix from one domain and the matrix helix of the neighboring domain. At sites 1 and 3, cardiolipin bridged adjacent domains through extensive engagement of each of its phosphates (Fig. 1*B*). At Site 2, a single phosphate of the cardiolipin molecule bridged the domains in 3 out of 4 replicates (Fig. 1*B*). This suggested at least two binding modalities for cardiolipin and that cardiolipin may engage SLC25A51 asymmetrically.

To determine whether the observed cardiolipin binding was required for the activity of SLC25A51, we expressed a genetically-encoded ratiometric fluorescent sensor for free NAD^+^ in the mitochondria of HeLa cells (1, 15, 29). Fig. 1*C* depicts the inverted fluorescence intensity of the sensor that is a readout of relative NAD^+^ levels. Acute ectopic expression of ^Flag^SLC25A51 resulted in increased steady-state levels of free mitochondrial NAD^+^ compared to empty vector, demonstrating that this assay could measure the activity of SLC25A51 of importing NAD^+^ in intact and respiring cells (Fig. 1*C*). We introduced point mutations to impair SLC25A51’s interaction with cardiolipin by blunting the electrostatic interactions at each binding sites (R82Q, R174Q, and R270Q). These sites were exterior to the transporter’s pore, distinct from the active site, and not obviously involved in any interhelical interactions. Mutation of all three sites, resulted in loss of SLC25A51 activity (Fig. 1*C*), indicating that cardiolipin is likely required for SLC25A51 activity.

### The matrix gate is comprised of a single salt bridge interaction

To study the control of SLC25A51 opening to the matrix, we examined all possible inter-helical interactions at the position where gated salt-bridge networks have been identified in the related MCF carriers (4, 6, 30). In SLC25A51, there were only two potential interhelical interactions: a putative hydrogen bond between Q52 and Q142, and a putative ionic salt-bridge between E139 and K236 (Fig. 2*A*). Time evolution of each interaction over 1000 ns in both AlphaFold2 and Swiss-Model models indicated that the hydrogen bond (Q52-Q142) was not formed (combined mean percent occupancy, m.p.o. 2% ± 0.7%) while the salt bridge interaction (E139-K236) was stable (combined m.p.o., 93% ± 3%) (Fig. 2*B* and *C*).

**Fig. 2.**
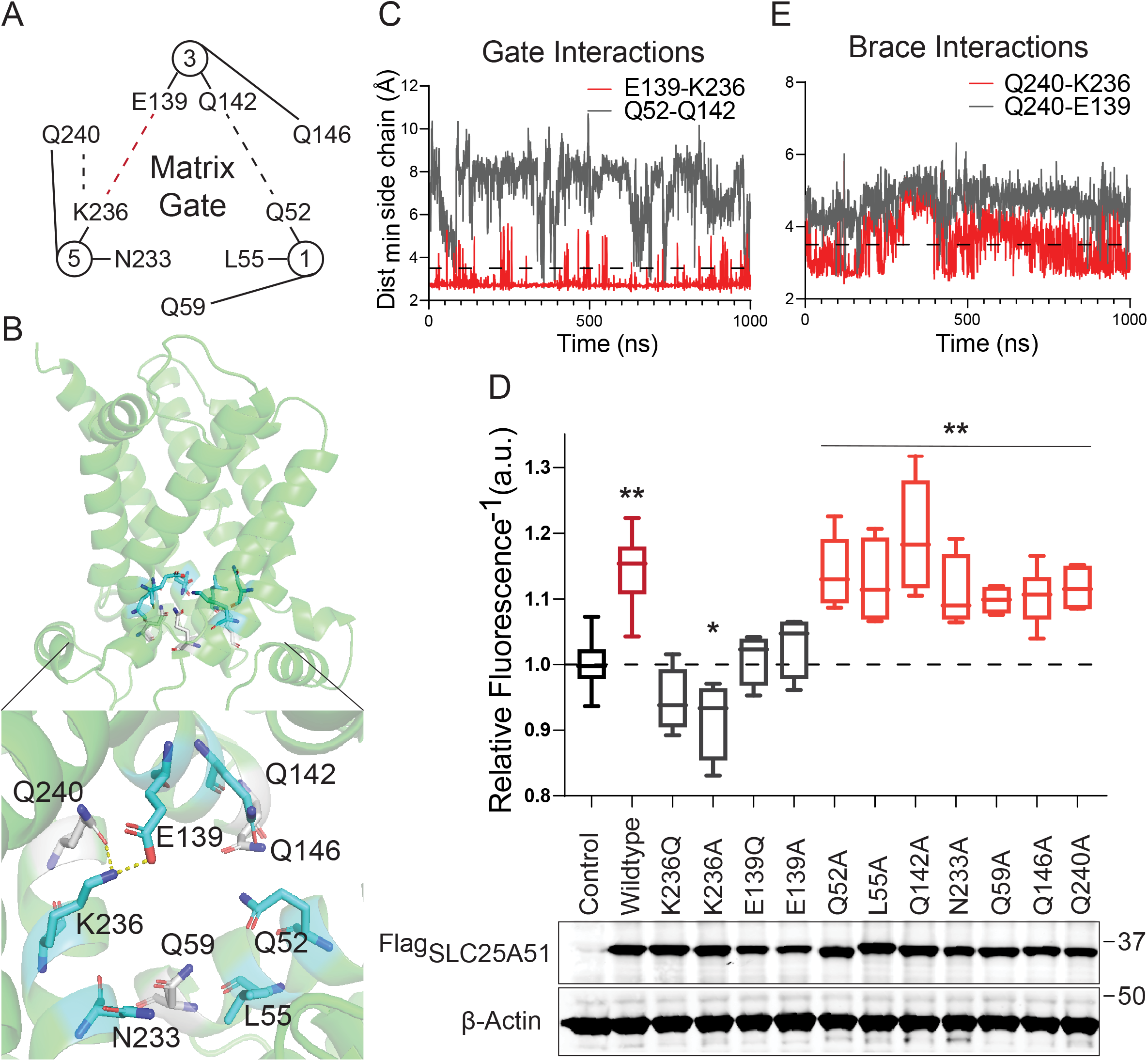
A single salt bridge stabilizes the NAD^+^-accessible, outward facing conformation. *(A)* Graphical representation of the matrix gate with dash lines indicating possible polar non-covalent interactions. Interactions were either indicated (red) or unsupported (grey) by data. *(B)* Relative position of the matrix gate in a cartoon representation of SLC25A51; tested glutamine brace residues (white) and interactions (dashed lines). *(C)* Time evolution of distance between indicated side chains for putative gate interactions. Interaction cutoff of 3.5 Å, dashed line. *(D)* Free mitochondrial NAD^+^ levels measured using a ratiometric biosensor in HeLa cells expressing Control empty vector (n=25), Wildtype ^Flag^SLC25A51 (n=25), and indicated mutants. Dashed horizontal line indicates baseline at 1. Data is mean ± SD, n = 4 - 6, ANOVA *p* <0.001 with post-hoc Dunnett’s test *\*p* < 0.05, *\*\*p* < 0.01. *(bottom)* Anti-Flag Western Blot from HeLa whole cell lysates express either empty vector or indicated ^Flag^SLC25A51 variants; β-actin, loading control. *(E)* Time evolution of distance between indicated side chains of putative brace interactions. Interaction cutoff at 3.5 Å, dashed line.

We used the mitochondrial NAD^+^ sensor assay to test whether these interactions were required for SLC25A51 activity. Loss of the gate is predicted to favor the inward-facing, matrix (m-state) conformation, which would not be amenable to NAD^+^ import. Expression of SLC25A51 variants with an ablated salt bridge (K236Q, K236A, E139Q, or E139A point mutations) were not able to increase steady-state levels of free mitochondrial NAD^+^ levels compared to empty vector control (Fig. 2*D*). This indicated that formation of the E139-K236 salt bridge was required for SLC25A51 activity. In line with the simulation results, disruption of the putative hydrogen bond with Q52A or Q142A did not have any major impact on SLC25A51 activity (Fig. 2*D*). Furthermore, conserved residues in the consensus matrix gate motif (PX[D/E]XX[K/R]XXQ)—including L55 and N233 whose equivalents were identified as critical in the yeast and plant mitochondrial carriers (31)—were not required for SLC25A51 activity in intact cells (Fig. 2*D*). Together, the modeling and the biochemical assays indicated that a single interhelical salt-bridge between E139 and K236 formed the matrix gate.

### Glutamine braces are not required for SLC25A51 activity

In many MCF carriers, glutamine braces play important roles in regulating the matrix gate (6). We observed an analogously positioned glutamine residue Q240 in SLC25A51 to putatively brace the identified E139-K236 salt-bridge (Fig. 2*A* and *B*). Modeling analyses was used to test the stability of either a Q240-E139 or Q240-K236 interaction. We did not observe consistent interactions in either model (Q240-E139 combined m.p.o. of 5% ± 3%; Q240-K236 combined m.p.o of 45% ± 16%) (Fig. 2*E*). In agreement, mutation of Q240A, or analogous glutamine residues Q59 and Q146 in adjacent domains, minimally impaired SLC25A51 activity (Fig. 2*D*). Together, our data demonstrated that the identified salt-bridge gate in SLC25A51 can function without stabilization from glutamine braces, distinctly unique from many other MCF carriers (6).

### Arginine cap residues contribute to regulating SLC25A51 activity

Cap residues at the matrix opening of the transporter have been proposed to help stabilize the c-state of MCF carriers by neutralizing c-terminal negative dipoles of converging odd-numbered helices or by forming stabilizing ionic interactions with matrix helices (32, 33). In SLC25A51’s modeling, we observed three arginine residues, R57, R238, and R140, positioned to putatively serve as cap residues (*SI Appendix*, Fig. S4*A*). While mutation of all three residues (R57Q, R140Q, and R238Q) destabilized the protein as observed by Western blot (*SI Appendix*, Fig. S4*B*), individual mutations were better tolerated and each partially impaired SLC25A51 activity (*SI Appendix*, Fig. S4*C*). In agreement, we observed stable interactions over time between R140 and the backbone of residue Q240 (m.p.o 95% ± 3% AlphaFold2 model, m.p.o. 80% ± 16% Swiss-Model), as well as between R238 and the backbone of residue Q59 (m.p.o. 70% ± 10%, Swiss-Model) (*SI Appendix*, Fig. S4*D-F*).

### NAD^+^ import can occur independently of a stable cytoplasmic gate

SLC25A51 is predicted to function via an alternating-access mechanism similarly to other MCF family members (4, 7). We thus generated an m-state model (MolProbity 94^th^ percentile)—using Swiss-Homology Modeling and the solved structure of *T. thermophila* ADP/ATP carrier (PDB ID: 6GCI) (25)—to help identify the cytoplasmic gate. Performed in triplicate (~1 μs / replicate, *SI Appendix*, Table S1), we observed consistent and stable formation of a putative ionic gate between K198 and E291 (m.p.o. 95% ± 5%) in an analogous plane to the identified cytoplasmic gate in the ADP/ATP carrier (*SI Appendix*, Fig. S5*A* and *B*). This suggested that SLC25A51 has single salt bridge gates on each side of its pore. Mutation of the cytoplasmic gate is expected to favor the c-state conformation. Interestingly, introduction of double mutation K198Q, E291Q did not impair SLC25A51 activity (*SI Appendix*, Fig. S5*C*) unlike previously observed for ADP/ATP carrier (11, 25). This indicated that a c-state conformation was sufficient for importing NAD^+^ and that a stable m-state was not required for continuous uptake of NAD^+^.

Putative tyrosine brace interactions were also observed but with lowered percent occupancy (E103-Y290 m.p.o. 68% ± 24%; R194-E291 m.p.o. 63% ± 8.5%) (*SI Appendix*, Fig. S5*A* and *B*). In agreement with the idea that a stable m-state was not essential, many of the disruptive mutations had minimal effect on SLC25A51 activity (*SI Appendix*, Fig. S5*C*). The exception was that R194A and R194Y blocked SLC25A51 activity, however mutation of its binding partner E291 did not result in any impairment, so any contribution from R194 was likely independent of its brace function. Together, the data indicated that cytoplasmic gate and brace features were not critical to SLC25A51’s import function.

### NAD^+^ is orientated by charge to bind at three conserved contact points

To study how SLC25A51 engages its ligand, we unbiasedly modeled the binding of NAD^+^ onto both c-state models. We used the Autodock 4.2 algorithm (34, 35) to generate a series of poses ordered by calculated binding energy scores. We chose the poses with the lowest binding energy scores from each model to continue with analyses using MD simulations.

With both models (*SI Appendix*, Table S1 Simulations 10-18), we observed a prompt (within 100 ns) and consistent positioning of the nicotinamide ring (Fig. 3*A*, *B* and Movies S2, S3). The central ligand-binding pore of SLC25A51 is characterized by positive charges arising from inward facing residues K91, R182 and R278 on the even-numbered helices, and a distinct negatively-charged region from acidic residue E132 on helix 3 (Fig. 3*A*). We observed in both models that the positively-charged nicotinamide ring of oxidized NAD^+^ consistently oriented away from the basic residues in the binding site and toward E132 (Fig. 3*A*, *B* and Movies S2, S3). Ablation of E132’s charge impaired SLC25A51 activity, but a E132D variant retained function (Fig. 3*C*). E132 is conserved across SLC25A51 orthologs (*SI Appendix*, Fig. S1*A*) (4).

**Fig. 3.**
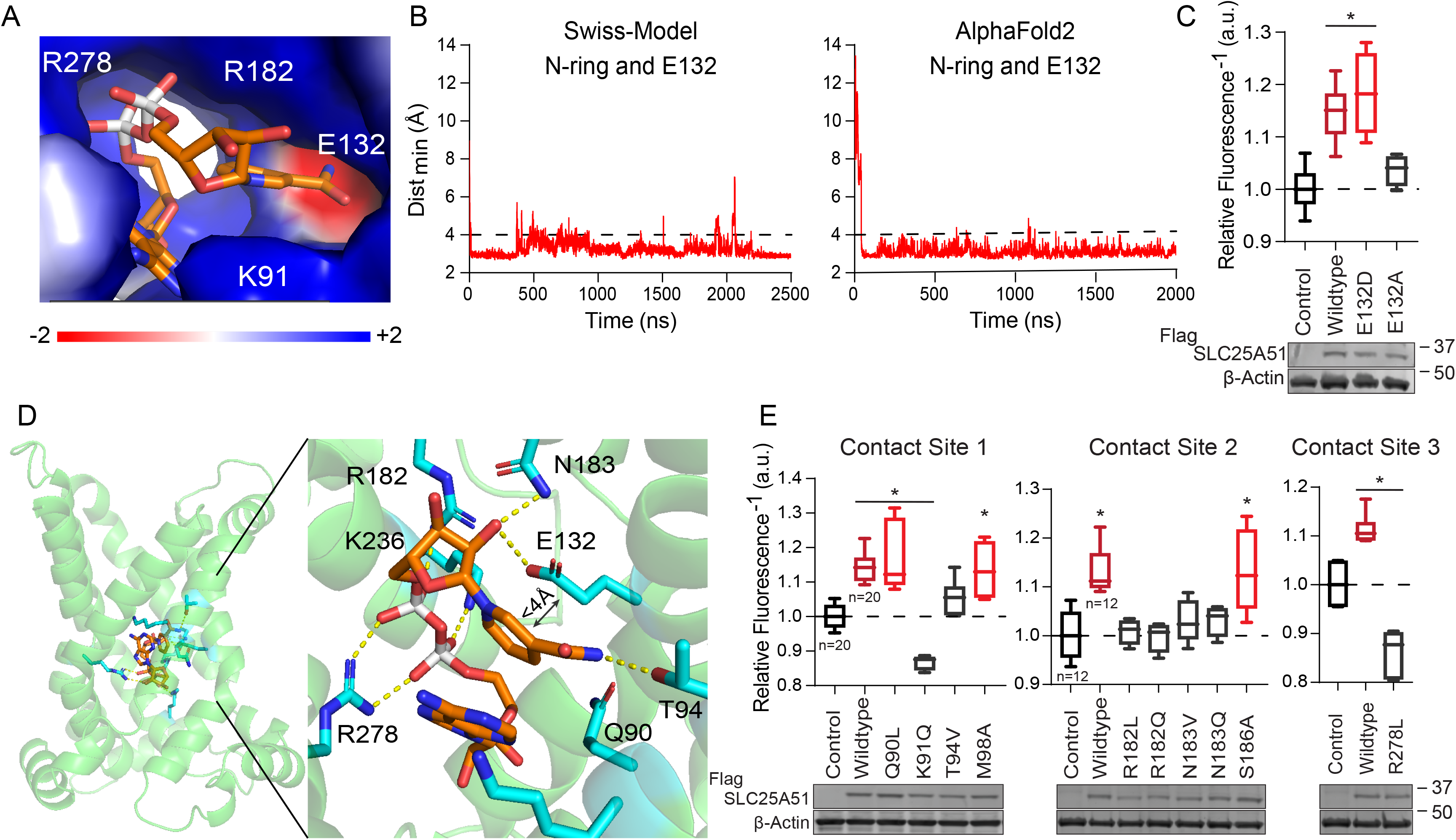
E132 and conserved contact sites comprise an NAD^+^ binding site. *(A)* Electrostatic environment of the substrate binding site (APBS as Pymol plugin). Representative positioning of NAD^+^ from simulation 17 at 1230 ns (*SI Appendix*, Table S1). *(B)* Minimal distance over time between the nicotinamide ring and E132 in simulations 17 (Swiss-Model) and 12 (AlphaFold2) (*SI Appendix*, Table S1). Interaction cutoff at 4 Å, dashed horizontal line. *(C)* Ratiometric sensor measurements of free mitochondrial NAD^+^ levels for ^Flag^SLC25A51 mutants E132D (n=4) and E132A (n=4) expressed in HeLa cells; Control empty vector (n=8) and Wildtype ^Flag^SLC25A51 (n=8). Dashed horizontal line indicates baseline at 1. Data is mean ± SD, ANOVA *p* < 0.001 with post-hoc Dunnett’s test *\*p* < 0.001. *(bottom)* Anti-Flag Western Blot; β-actin, loading control. *(D)* Bound NAD^+^ (orange) in the ligand binding pocket is shown in both side and enlarged views. Non-covalent interactions < 3.5 Å, dashed yellow lines. *(E)* Free mitochondrial NAD^+^ levels measured using a ratiometric biosensor in HeLa cells that expressed Control empty vector, Wildtype ^Flag^SLC25A51, or mutant variants as indicated. Dashed horizontal line indicates baseline at 1; data is mean ± SD, n = 4-7, unless otherwise indicated, ANOVA *p* < 0.001, post-hoc Dunnett’s test *\*p* < 0.001. *(bottom)* Anti-Flag Western Blot; β-actin, loading control.

Following positioning of the nicotinamide ring, we observed the nicotinamide ribose (NR) moiety to form the following consistent contacts (in 8 out of 9 simulations with both Swiss-Model and AlphaFold2 models) (*SI Appendix*, Table S1). Borrowing from the nomenclature used to describe the ADP/ATP binding pocket (7, 36, 37), we observed that residue T94 in contact site 1 and N183 in contact site 2 stabilized the NR moiety through specific hydrogen bonds with its carboxamide group and ribose (2’ and 3’ OH) respectively (Fig. 3*D*). R278 (contact site 3), R182, and K91 engaged the negative charges from the phosphates of NAD^+^ (Fig. 3*D* and Movies S2, S3).

We mutated and tested the requirement of every conserved residue in the three contact sites using the sensor assay (Fig. 3*E* and *SI Appendix*, Fig. S1) (4, 6). For contact site 1, we found that SLC25A51 variants with a T94V or K91Q mutation impaired SLC25A51 import with minimal effects on protein stability. Mutating adjacent contact site residues (Q90 and M98) that were not observed to participate in binding of the NR moiety minimally impaired SLC25A51 despite being conserved residues (Fig. 3*E*). Similarly, we confirmed that R182 and N183 were required for site 2 but not neighboring residue S186, and we confirmed that R278 was required for site 3 (Fig. 3*E*).

### NAD^+^ directly engaged with the gate in SLC25A51

While modeling the binding of NAD^+^, we unexpectedly observed two simulations wherein the ligand traversed deeper into the pore (Movie S4 and *SI Appendix*, Table S1 simulations 16 and 17). To determine whether an extended simulation could provide additional insights, we continued with a pose that originated from the unbiased docking with the Swiss-Homology model but had scored among lowest binding energy scores (*SI Appendix*, Table S1 simulations 19-22). This pose positioned the nicotinamide ring deeper in the pocket.

In all four replicate simulations 19-22, we observed channeling of the nicotinamide ring from E132 to E139 (Fig. 4*A*, *B* and Movie S4). In both model structures residue E139, of the salt-bridge gate, was positioned below E132 on the same helix 3. We also observed that as the nicotinamide ring interacted with E139 (85%, 48%, 69% and 93% occupancy), the phosphates on NAD^+^ could interact with K236, the other half of the salt bridge. Resultingly, formation of the E139-K236 salt bridge was negatively correlated with the NAD^+^ phosphate-K236 interaction (r^2^ = −0.192, −0.045, −0.681 and −0.336 per replicate, *p* < 0.001), as well as with the nicotinamide ring-E139 interaction (r^2^ = −0.298, −0.053, −0.091 and −0.342 calculated per replicate, *p* < 0.001) (Fig. 4*C*, *D* and Movie S5).

**Fig. 4.**
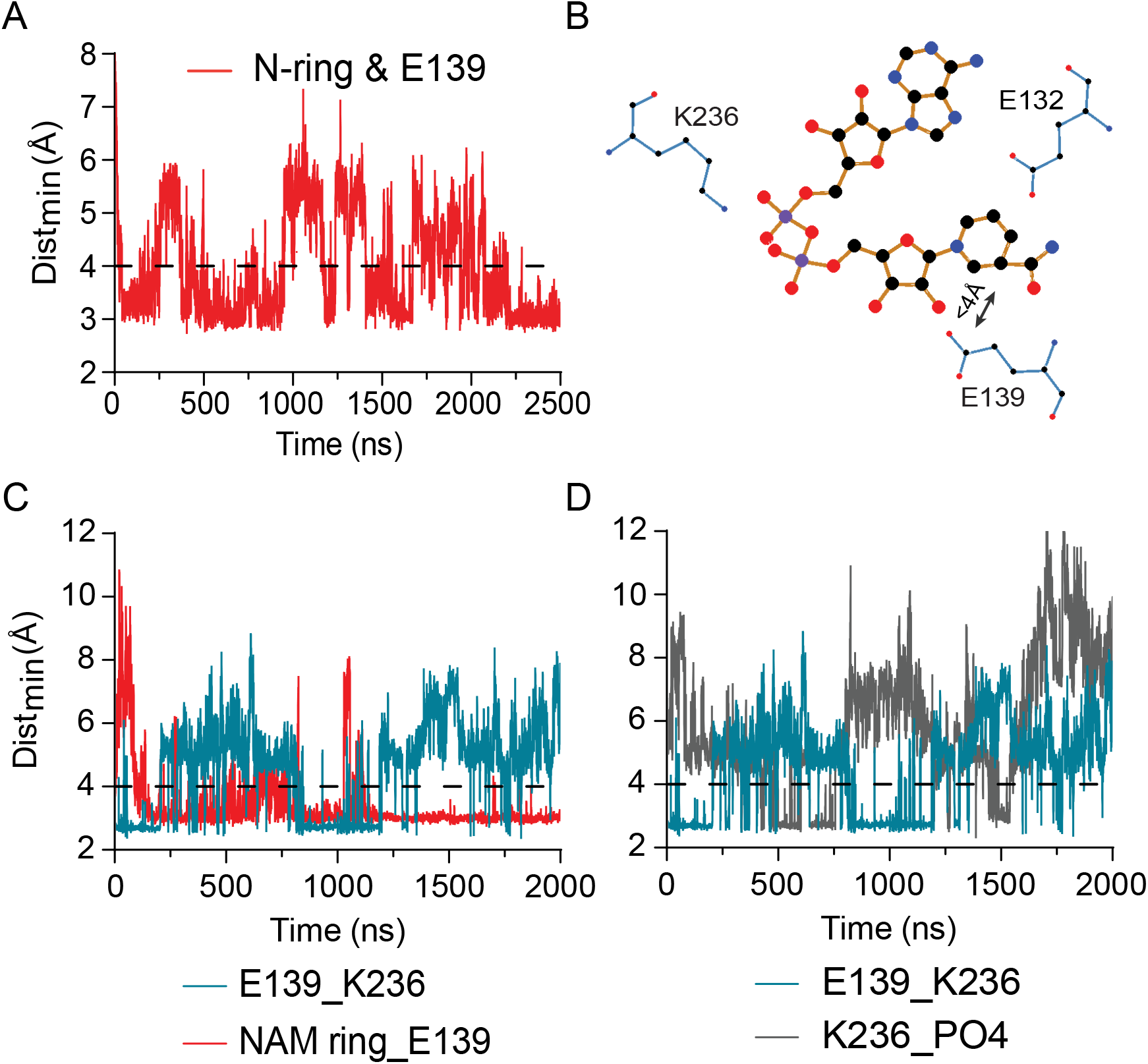
Substrate induced disruption of the matrix gate. *(A)* Minimum distance between the nicotinamide ring and E139 over time, simulation 17 (*SI Appendix*, Table S1). *(B)* 2-dimentional projection (Ligplot+) of NAD^+^ and its interaction with E132 and the E139-K236 salt bridge. *(C)* Time evolutions of the E139-K236 salt bridge (teal) compared to the nicotinamide ring and E139 interaction (red); simulation 19 (*SI Appendix*, Table S1) *(D)* Time evolutions of the E139-K236 salt bridge (teal) compared to the phosphates in NAD^+^ and K236 (grey); simulation 19 (*SI Appendix*, Table S1)

## Discussion

The molecular characterization of mitochondrial carriers has been a persistent challenge with crystallographic structures currently available only for the bovine and fungal ADP/ATP carriers (5, 8, 9, 25, 38). SLC25A51’s lack of close homology to the previously identified yeast and plant NAD^+^ transporters delayed its de-orphanization as a mammalian mitochondrial NAD^+^ transporter (*SI Appendix*, Fig. S1). Using a combination of modeling, simulations, and biochemical assays, we have gained insight into SLC25A51’s binding of NAD^+^, its dependency on cardiolipin for function, and a ligand induced mechanism that disrupted the matrix gate.

A unique aspect in NAD^+^’s binding to SLC25A51 was the initial orientation of its oxidized nicotinamide ring via electrostatic interactions. We observed that the delocalized positive charge on the ring was drawn to the negative surface on helix 3 formed by E132 and simultaneously repelled from basic residues K91, R182, and R278 in the same plane (Fig. 3*A*, Movie S2 and S3). This positioned NAD^+^ such that the nicotinamide ribose moiety engaged contact sites 1 and 2 consistently, and the negative charges on NAD^+^’s phosphate groups were satisfied and engaged with the positively charged sidechains in the pore (e.g., R278 in contact site 3) (Fig. 3*D*). In addition to the negative charge on E132, we confirmed that the residues comprising the identified contact sites were required for SLC25A51 activity (Fig. 3*C* and *E*). The data was consistent with previous docking and mutational analyses (2, 4). We observed that all three contact sites in SLC25A51’s pore must be engaged to orient the ligand such that it can travel deeper into the pore. This may explain why nicotinamide, or its mononucleotide were unable to efficiently compete with NAD^+^ for transport (1, 2).

Positioned below NAD^+^’s initial binding site in SLC25A51 was a single interhelical salt-bridge that embodied the matrix gate (Fig. 2). This salt-bridge interaction was the only consistent interhelical bond formed within SLC25A51’s PX[D/E]XX[K/R] motifs, and it was required for activity (Fig. 2*D*) (6). Following the initial binding of NAD^+^, the nicotinamide ring was channeled to the residue E139, which is located directly below E132 and comprises half of the matrix gate (Movie S4). The negatively charged phosphate groups on NAD^+^ in turn engaged the other half of the gate, residue K236, and together this disrupted an otherwise stable gate interaction (Fig. 4 and Movie S5). However, it will still need to be determined whether this interaction can proceed to an occluded state or to the transport of the molecule. An analogous mechanism was proposed for the ADP/ATP carrier involving interference of the gate via the N6 atom of the adenine moiety on the nucleotide or the guanidino NH1 or NH2 of a conserved binding site arginine residue (36). We found additional features in SLC25A51 such as an arginine-cap that functioned to further stabilize the matrix gate (*SI Appendix*, Fig. S4), but we also identified that the c-state was sufficient for continuous import of NAD^+^ when the cytoplasmic gate was weakened indicating that its relatively weak gate could be readily opened as suggested previously (*SI Appendix*, Fig. S5) (7).

NAD^+^’s interaction with the matrix gate depended on the positive charge of its nicotinamide ring being positioned close to the acidic residue of the gate. This implies that without this positive charge, even with ideal positioning of the ring, the transport process may be halted. This model is consistent with the electrostatic funneling observed with ADP/ATP carriers (39–44). Moreover, previous studies have found that AMP, dUTP, NADH, GDP and GTP could bind to the ADP/ATP carrier, suggesting that the dynamics of transport in cells involves an additional differentiation step to discriminate between bound molecules (45, 46). Our substrate-induced model may explain why NAD^+^ can be transported at micromolar concentrations but that NADH can compete for SLC25A51 uptake at millimolar concentrations (2). Apart from an oxidized nicotinamide ring, NADH is structurally identical and likely can engage all binding sites in SLC25A51. Nevertheless, cytosolic ratios—which due to non-discriminating porin channels in the outer membrane—represent what is experienced by SLC25A51 in cells. Free NADH is approximately 100 to 1000-fold less abundant than free NAD^+^ (47–50). In this scenario the relatively subtle E132 may be enough to create a molecular discrimination based on electrostatic charge. Additionally, the overall charge of an NADH or an NAD^+^ molecule would be differentially masked in the positive pore of SLC25A51 (6). With an extra positive charge from its nicotinamide ring, NAD^+^ bound to SLC25A51 retains a net positive charge in the binding site is likely to be favored by the negative potential in the matrix, compared to the neutral charge of bound NADH in the same site (6).

To transport a ligand such as NAD^+^ with an asymmetric structure, one may either expect the pore to be matched in asymmetrical shape or to have flexibility. We found evidence suggesting that SLC25A51 required cardiolipin for its activity (Fig. 1). Cardiolipin has been found to support MCF activities in a variety of ways (28, 51–53). The data supporting cardiolipin recruitment at specific sites was consistent and collected unbiasedly. We observed two modalities of cardiolipin binding, and site 2 cardiolipin tended to bind non-canonically using only one of its phosphates. A potential consequence is that this would asymmetrically impact the pore in a dynamic manner that is potentially regulatable.

Overall, this study provided unique insights into the binding and transport mechanism of an NAD^+^ carrier. It is a launching point for future characterization and structural studies to answer remaining questions such as what controls the directionality of NAD^+^ import, whether SLC25A51 requires an antiport molecule akin to other carriers, and to elucidate the full dynamics of the SLC25A51 transporter. A structural understanding will accelerate the development of modulators targeting SLC25A51 in various diseases.

## Material and Methods

Detailed material and methods are provided in the Supporting Information Appendix.

## Competing Interests

The authors declare no competing interests

## Acknowledgements

We thank Akhilesh Paspureddi and Dr. Ron Elber for valuable discussions and help with simulation analyses throughout this study. Supported by the Texas Advanced Computing Center, NIH DP2 GM126897, the Pew Charitable Trust, and CPRIT RP210079.

## Supporting Information

### Materials and Methods

#### Multiple Sequence Alignment

Multiple Sequence Alignment using Cobalt sequence alignment tool of representative nucleotide carriers *Hs*SLC25A51 (UniProt: Q9H1U9), *Bt*SLC25A4 (UniProt: P02722, PDB ID: 1OKC), *Tt*ADT (UniProt: G2QNH0, PDB ID: 6GCI), *Sc*Ndt1 (UniProt: P40556) and *At*Ndt1 (UniProt: O22261) (1). The sequence alignment was viewed and analysed using Unipro UGENE (2).

#### SLC25A51 Models

The homology models of cytoplasmic open state of apo-SLC25A51 (27-297) and apo-SLC25A4 were generated using the Swiss-Model server with bovine SLC25A4 crystal structure as the template (PDB ID: 1OKC) (3, 4). Bovine SLC25A4 was chosen as it has a high-resolution crystal structure available (2.2 Å) and its GMQE score of 0.55 was the highest for SLC25A51 amongst all the c-state MCF carrier structures available. The homology model of mitochondrial matrix open state of apo-SLC25A51 (26-297) was generated using the Swiss-Model server with a thermophilic fungal ADP/ATP Carrier crystal structure as the template (PDB ID: 6GCI) (5). We also used AlphaFold2 generated cytoplasmic open state model for our study (6). The SLC25A51 models quality estimation was performed using MolProbity, pLDDT scores and QMEANDisCo analysis (6–8).

#### Molecular Dynamics Simulation system setup

CharmmGUI webserver was used to build the starting coordinates of our system for MD simulations with a total dimension of 85 Å × 85 Å × 100 Å (9, 10). The c-state Swiss-Model (apo–state or NAD^+^ docked), m-state Swiss-Model and apo-c-state AlphaFold2 SLC25A51 were embedded in lipid bilayer systems with phosphatidylcholine (POPC), phosphatidylethanolamine (POPE) and cardiolipin (TLCL2) in a ratio 2:3:2 as experimentally determined previously (11). We chose TLCL2 cardiolipin with a negative 2 charge that agrees with the recent studies on ionization properties of the cardiolipin molecules (12, 13). The protein and the lipid bilayer were solvated with TIP3P water molecules extending up to 22.5 Å on both sides of the membrane. The system also contains neutralizing K^+^ and Cl^−^ ions to make the final salt concentration 150 mM. The NAD^+^ was docked on the final coordinate of SLC25A51 after 1 μs simulation of AlphaFold2 SLC25A51 model which was embedded in a lipid bilayer membrane of POPC lipids with all the other parameters same as the other models described above.

#### Molecular Dynamics Simulation

The simulations were performed with Gromacs (version 2022.1) using an all atom Charmm36m force field for protein, lipids, ions and ligands with periodic boundary conditions (14, 15). We performed energy minimization of the system in 5000 steps of steepest descent with position restraints on protein and NAD^+^ while lipids, water and ions were free to equilibrate. The system was equilibrated in a multi-step procedure with decreasing positional and dihedral constraints with each step with following parameters: Two steps of 0.125 ns NVT simulation at 310.15 K, one step of 0.125 ns NPT simulation at 310.15 K and 1 bar and three steps of 0.5 ns NPT simulation at 310.15 K and 1 bar. MD simulations were run for different lengths of time using leap-frog algorithm as an integrator with 2 fs time step. Particle Mesh Ewald was used for non-bonded long-range interactions with a 12 Å cutoff and Leonard-Jones potential cutoff was 12 Å. Temperature was maintained at 310.15 K using Nose-Hoover thermostat with a 1 ps coupling parameters and pressure at 1 bar using Parrinello-Rahman barostat with 5 ps coupling and 4.5 × 10^−5^ bar^−1^ compressibility.

#### Trajectory analysis

The analysis was done using in-built tools available in the gromacs package. The coordinates were viewed, and structural graphics and videos were prepared using Pymol. The cutoff distance was 3.5 Å for hydrogen bonds and for salt bridges while 4 Å for distance between any atom of Nicotinamide ring of NAD^+^ and side chain of amino acid. The mean percentage occupancy (mpo) represents the percentage of simulation time where the distance between the atoms was under the cut-off distance. Time evolution of distances and RMSD is shown as moving averages of 5 values on each side. The videos are made with every 10^th^ ns frame of the trajectories

#### APBS Electrostatics

APBS electrostatics was performed using Pymol plugin. Molecule was prepared using pdb2pqr method with 0.5 grid spacing and the electrostatic potential was projected to Connolly Surface in the −2 to +2 range.

#### Cell Culture

HeLa (ATCC: CCL-2) cells were cultured in Dulbecco’s modified Eagle’s medium (DMEM) containing 4.5 gL^−1^ glucose, 1 mM sodium pyruvate and 4 mM L-glutamine supplemented with 10% fetal bovine serum, 1× penicillin–streptomycin and 12.5 mM HEPES.

#### Cloning and mitochondrial NAD^+^ measurements

Q5 site directed mutagenesis kit from NEB was used to make mutations in pCMV-Flag-HA-SLC25A51(1-297)-IRES-puro plasmid described previously (16). The plasmids were transiently transfected in HeLa ^mito^cpVenus and HeLa ^mito^Sensor cells using polyethylenimine (PEI, 1 mg/mL) in a PEI:DNA ratio of 5:1. Relative NAD^+^ steady state measurements were taken as described in our previous work in detail (17). The relative fluorescence depicted in the figures represents the NAD^+^ sensor fluorescence normalized to the CpVenus control.

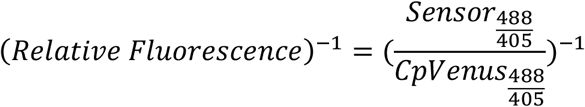

#### Western Blotting

Cells were lysed with 2x Laemmli sample buffer containing DTT. Lysates from 125000 cells were resolved using NuPAGE™ 10% or 4–12% Bis-Tris protein gels (Invitrogen) and were transferred to Bio-Rad 0.45 μm nitrocellulose membrane. The nitrocellulose membrane was blocked with 5% BSA in pH 7.4 Tris-buffered saline with 0.1% (v/v) Tween 20 (TBST). The antibodies anti-Flag M2 (Sigma, F1804, 1:2,000), anti-β-Actin (Sigma, A2228, 1:3000) and anti-α-Tubulin (Sigma T9026, 1:3000) were used for immunoblotting and were prepared in TBST with 1% BSA.

#### Statistical Analysis and Figure Graphics

The results are presented as mean ± SD in box and whisker graphical format. The data was analyzed, and figures were prepared using GraphPad Prism version 9, Microsoft Excel and adobe illustrator. One-way Anova was used for comparing biosensor measurements between 3 or more samples and multiple comparison was done using the Dunnett’s test. P-values under 0.05 were considered significant and the figure legend for each figure describes the significance criteria and number of replicates in that figure.

## Movie Legends

**Movie S1 (separate file)**: Cardiolipin molecules bind specifically to three conserved binding sites on SLC25A51 displacing other lipids in 1 μs of simulation 2 (*SI Appendix*, Table S1). SLC25A51 is shown from the matrix side as cartoon and cardiolipin molecules in sticks representation. Other lipids are not shown for clarity.

**Movie S2 (separate file)**: A GIF with 4 frames of the first 100 ns of simulation 17 (*SI Appendix*, Table S1) shows the reorientation of NAD+ with the positively charged nicotinamide ring near E132 and adoption of the binding site described in Fig. 3. Basic residues from contact sites K91, R182 and R278 are shown in cyan, N183 in wheat and E132 residue on Helix 3 is shown in red color.

**Movie S3 (separate file)**: First 1.5 μs of simulation 12 of AlphaFold2 model (*SI Appendix*, Table S1) shows the reorientation of NAD+. The representation of structures and coloring scheme is same as used in Movie S2.

**Movie S4 (separate file)**: Nicotinamide ring is guided by E132 (red, above) deeper into the binding pore and transiently interacts with E139 (red, below) salt bridge residue. The movie represents 2.5 μs of simulation 17 (*SI Appendix*, Table S1).

**Movie S5 (separate file)**: The NAD+ molecule disrupts the salt bridge residue with direct interaction of nicotinamide ring with E139 and NAD+ phosphate with K236. The movie represents 2 μs of simulation 19 (*SI Appendix*, Table S1) with NAD+ docked closer to nicotinamide ring.

**Fig. S1.**
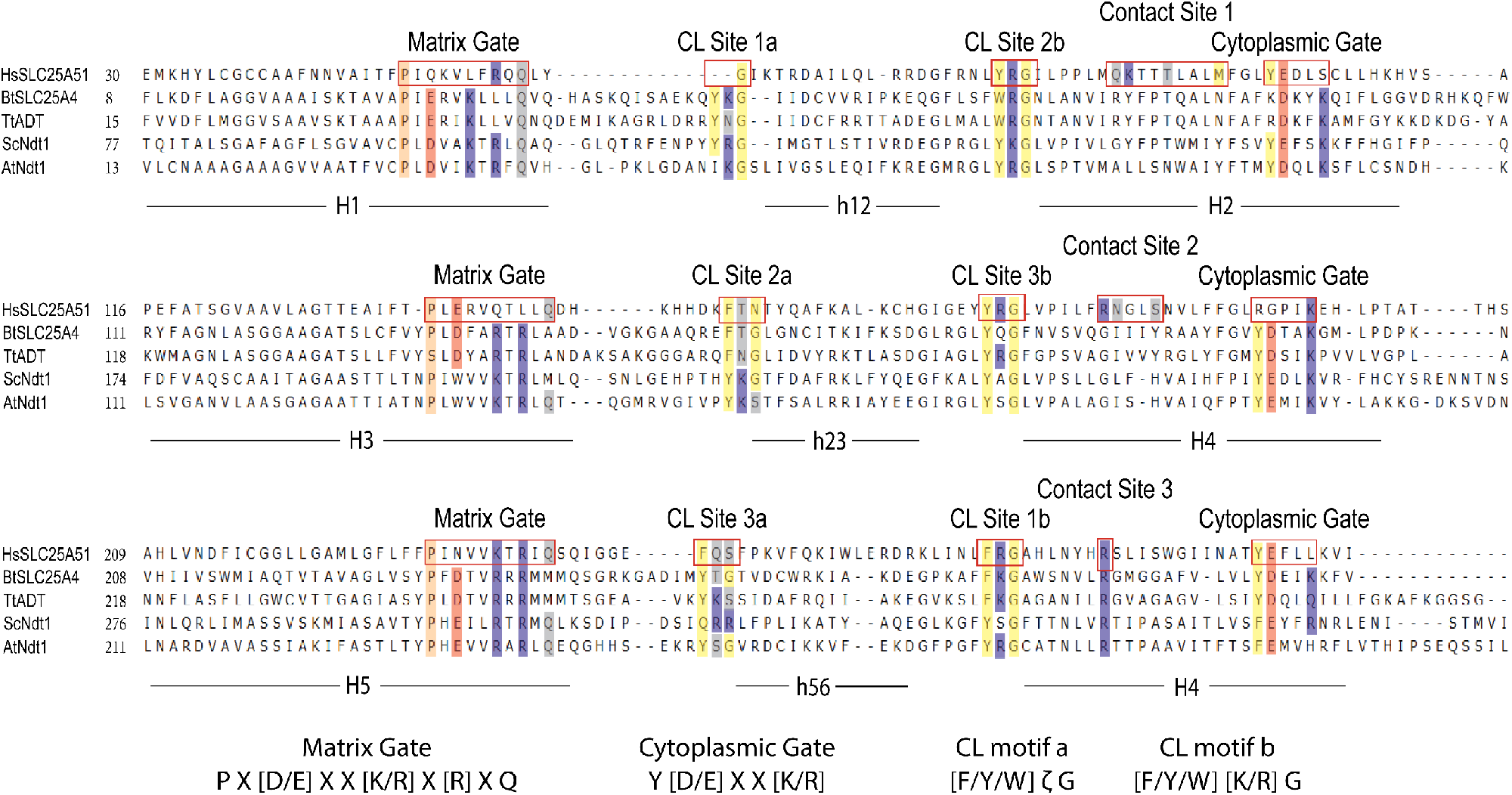
Multiple sequence alignment of representative nucleotide carriers. Alignment of amino acid sequences for *Hs*SLC25A51, *Bt*SLC25A4, *Tt*ADT, *Sc*Ndt1 and *At*Ndt1. Highlighted hydrophobic residues are depicted in yellow, hydrophilic in grey, negative residues in red, positive residues in blue and proline kink residues in peach.

**Fig. S2.**
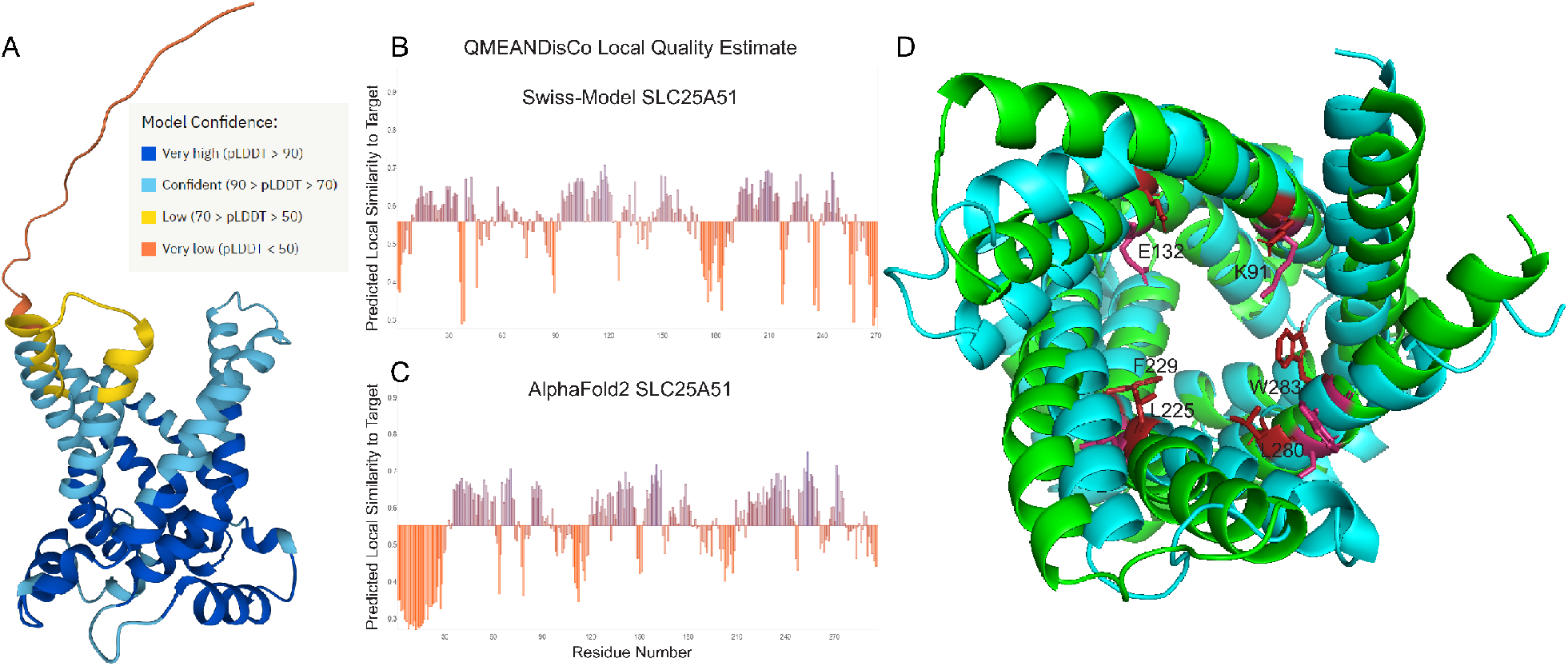
Quality Estimation and difference between Swiss-Model and AlphaFold2 structures. *(A)* Model confidence for AlphaFold2 structure of HsSLC25A51. *(B and C)* QMEANDisCo Local Quality Estimate shows higher confidence for transmembrane helices and low for flexible loops. (D) Alignment of Swiss-Model (Cyan) and AlphaFold2 (Green) structures. Key residues that are differentially positioned in the binding site are highlighted.

**Fig. S3.**
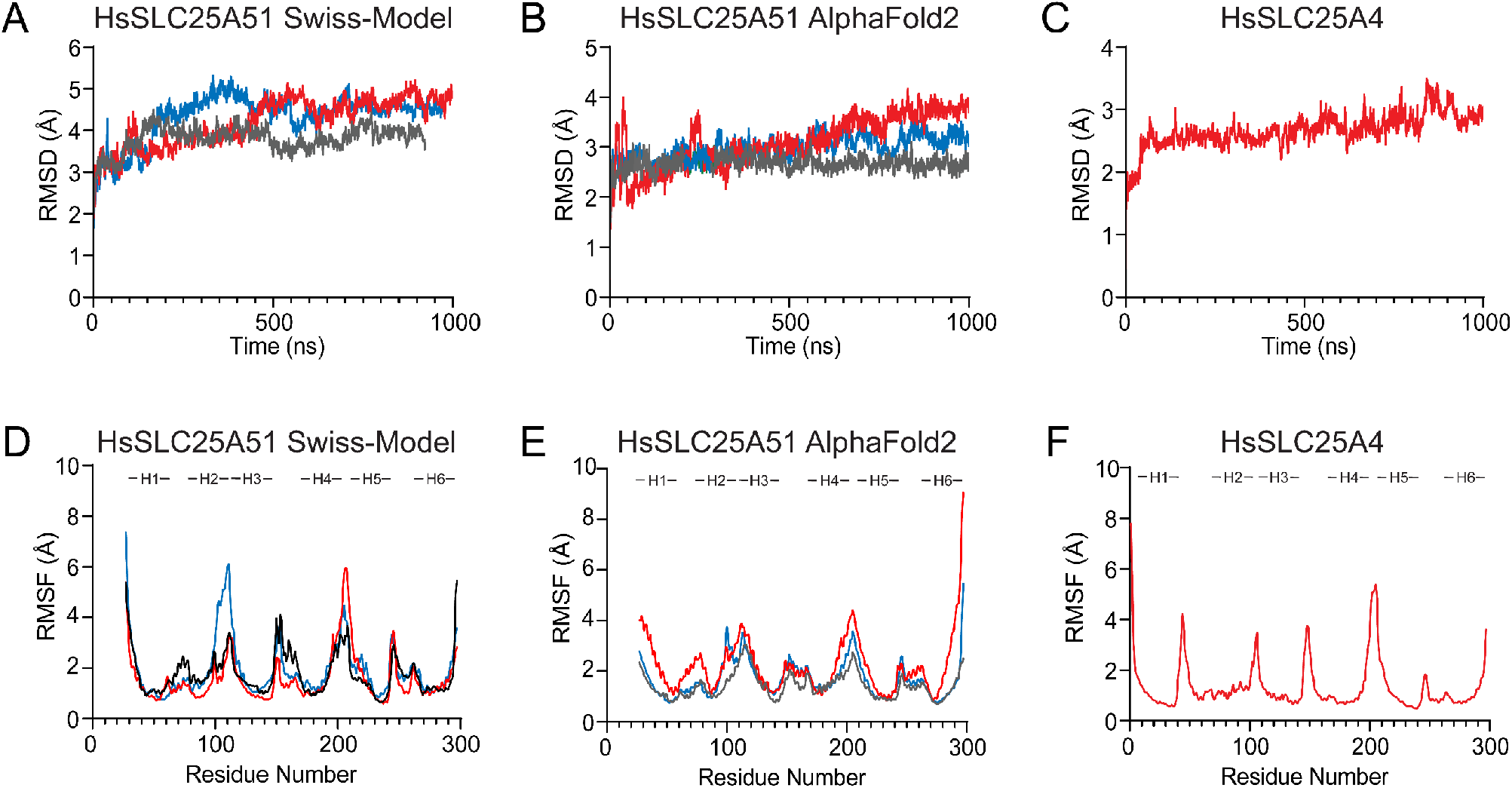
Root Mean Square Deviation and Root Mean Square Fluctuation of the simulations. RMSD of backbone atoms over time for *(A)* apo Swiss-Model (27-297 residues, n=3), *(B)* apo AlphaFold2 (27-297 residues, n=3) and *(C)* apo SLC25A4 (n=1) structures. RMSF of each residue for *(D)* apo Swiss-Model (27-297 residues, n=3), *(E)* apo AlphaFold2 (27-297 residues, n=3) and *(F)* apo SLC25A4 (n=1) structures.

**Fig. S4.**
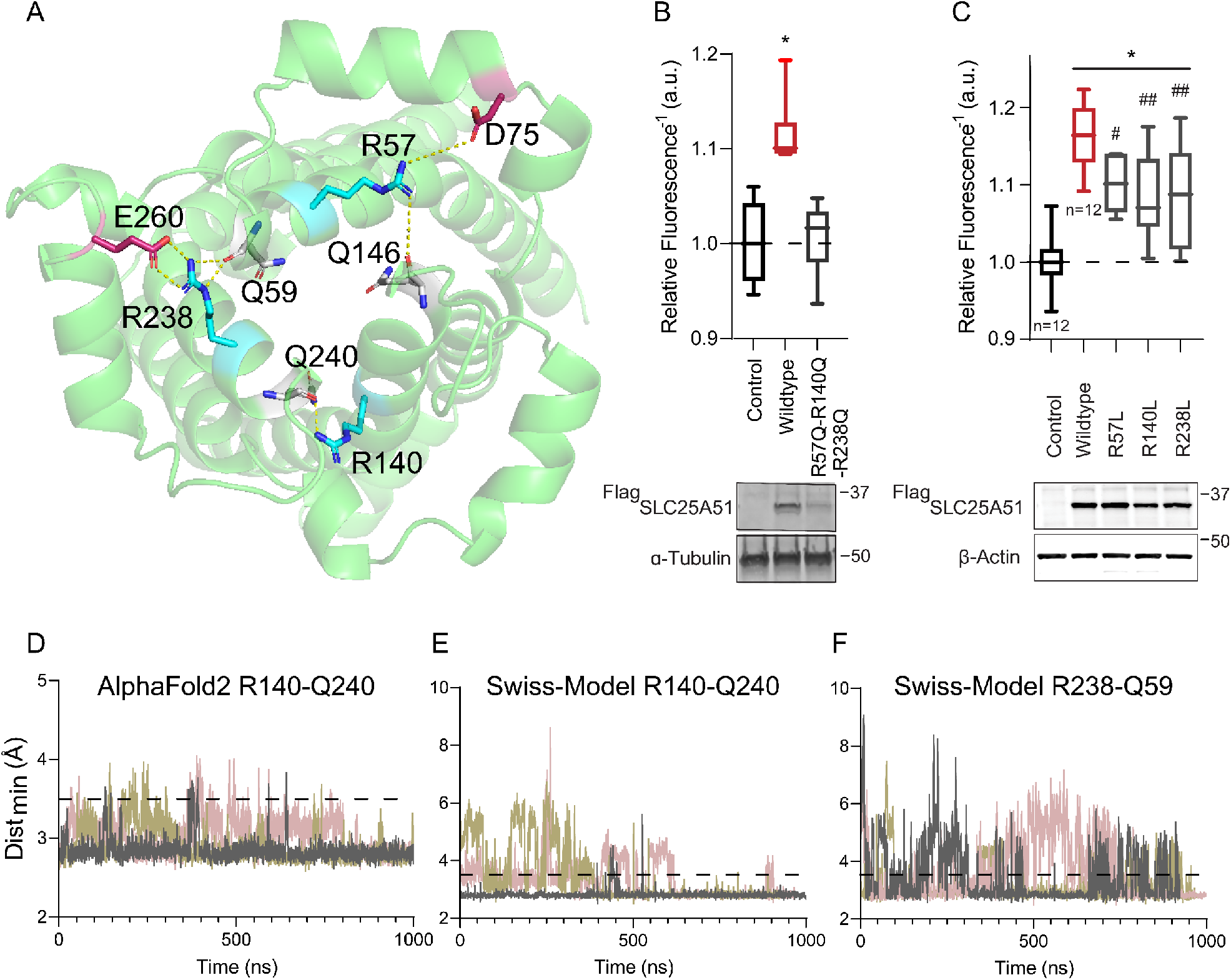
Cap residues are important for SLC25A51 function. *(A)* Relative positions of cap residues and their interactions (dashed lines) in a cartoon representation of SLC25A51. *(B and C)* Free mitochondrial NAD+ levels measured using ratiometric biosensor in HeLa cells that expressed Control empty vector, Wildtype ^Flag^SLC25A51, or mutant variants as indicated. Dashed horizontal line indicates baseline at 1; data is mean ± SD, n=6 unless otherwise indicated, ANOVA *p* <0.001, post-hoc Dunnett’s test compared to Control empty vector (*) and compared to Wildtype ^Flag^SLC25A51 (#), \**p* <0.01, #*p* <0.05 *and* ##*p* <0.01. *(bottom)* Anti-Flag Western Blot; Loading control as indicated. *(D-F)* Time evolution of distance between Arg side chains and Gln backbone forming the cap interactions for the indicated models. Interaction cutoff at 3.5 Å, dashed horizontal line.

**Fig. S5.**
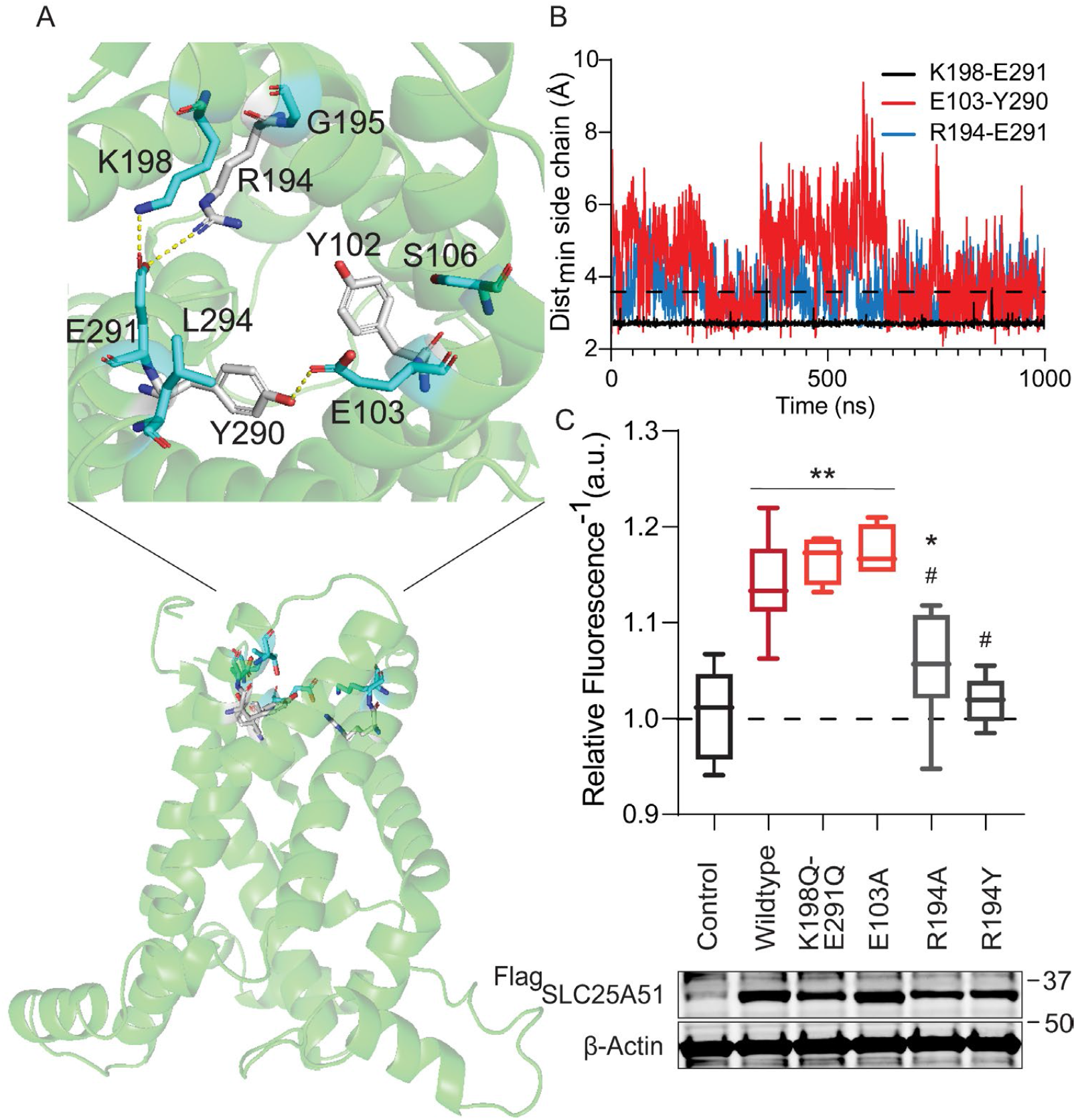
Cytoplasmic gate is not needed for NAD^+^ import by SLC25A51. *(A)* Relative position of the cytoplasmic gate in a cartoon representation of SLC25A51. *(B)* Time evolution of distance between indicated side chains for putative gate interactions. Interaction cutoff at 3.5 Å, dashed horizontal line. *(C)* Free mitochondrial NAD+ levels measured using ratiometric biosensor in HeLa cells that expressed Control empty vector (n=15), Wildtype ^Flag^SLC25A51 (n=15), ^Flag^SLC25A51 harboring K198Q and E291Q mutations (n=4) and mutants E103A (n=4), R194A (n=7) and R194Y (n=5). Dashed horizontal line indicates baseline at 1; data is mean ± SD, ANOVA *p* <0.0001, post-hoc Dunnett’s test compared to Control empty vector (*) and compared to Wildtype ^Flag^SLC25A51 (#), \**p* <0.05, \*\**p* <0.0001 *and* #*p* <0.001. *(bottom)* Anti-Flag Western Blot; β-actin, loading control.

**Table S1.**
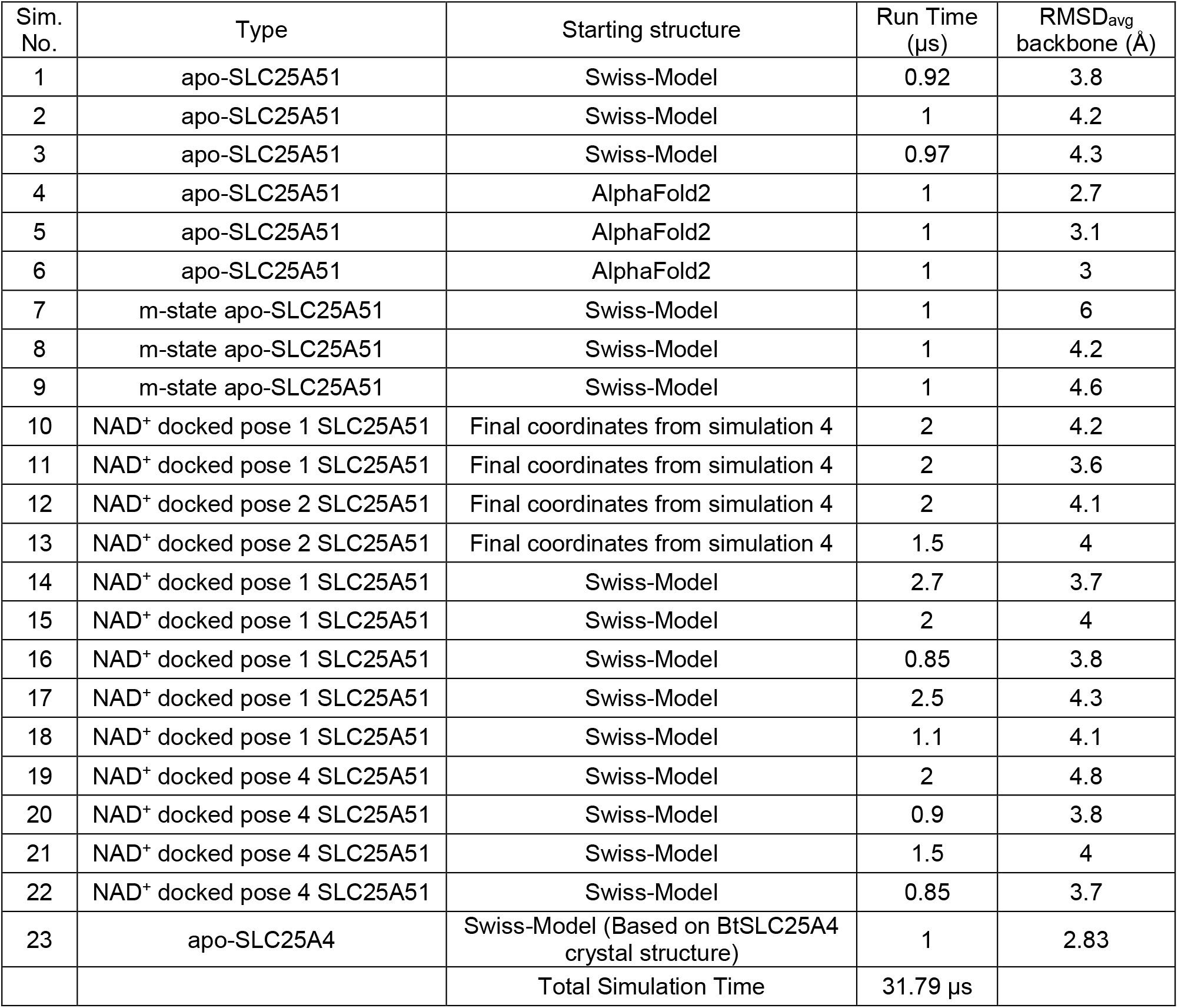
A summary of all the simulations performed in this study.

| Sim. No. | Type | Starting structure | Run Time (μs) | RMSD<sub>avg</sub> backbone (Å) |
| --- | --- | --- | --- | --- |
| 1 | apo-SLC25A51 | Swiss-Model | 0.92 | 3.8 |
| 2 | apo-SLC25A51 | Swiss-Model | 1 | 4.2 |
| 3 | apo-SLC25A51 | Swiss-Model | 0.97 | 4.3 |
| 4 | apo-SLC25A51 | AlphaFold2 | 1 | 2.7 |
| 5 | apo-SLC25A51 | AlphaFold2 | 1 | 3.1 |
| 6 | apo-SLC25A51 | AlphaFold2 | 1 | 3 |
| 7 | m-state apo-SLC25A51 | Swiss-Model | 1 | 6 |
| 8 | m-state apo-SLC25A51 | Swiss-Model | 1 | 4.2 |
| 9 | m-state apo-SLC25A51 | Swiss-Model | 1 | 4.6 |
| 10 | NAD<sup>+</sup> docked pose 1 SLC25A51 | Final coordinates from simulation 4 | 2 | 4.2 |
| 11 | NAD<sup>+</sup> docked pose 1 SLC25A51 | Final coordinates from simulation 4 | 2 | 3.6 |
| 12 | NAD<sup>+</sup> docked pose 2 SLC25A51 | Final coordinates from simulation 4 | 2 | 4.1 |
| 13 | NAD<sup>+</sup> docked pose 2 SLC25A51 | Final coordinates from simulation 4 | 1.5 | 4 |
| 14 | NAD<sup>+</sup> docked pose 1 SLC25A51 | Swiss-Model | 2.7 | 3.7 |
| 15 | NAD<sup>+</sup> docked pose 1 SLC25A51 | Swiss-Model | 2 | 4 |
| 16 | NAD<sup>+</sup> docked pose 1 SLC25A51 | Swiss-Model | 0.85 | 3.8 |
| 17 | NAD<sup>+</sup> docked pose 1 SLC25A51 | Swiss-Model | 2.5 | 4.3 |
| 18 | NAD<sup>+</sup> docked pose 1 SLC25A51 | Swiss-Model | 1.1 | 4.1 |
| 19 | NAD<sup>+</sup> docked pose 4 SLC25A51 | Swiss-Model | 2 | 4.8 |
| 20 | NAD<sup>+</sup> docked pose 4 SLC25A51 | Swiss-Model | 0.9 | 3.8 |
| 21 | NAD<sup>+</sup> docked pose 4 SLC25A51 | Swiss-Model | 1.5 | 4 |
| 22 | NAD<sup>+</sup> docked pose 4 SLC25A51 | Swiss-Model | 0.85 | 3.7 |
| 23 | apo-SLC25A4 | Swiss-Model (Based on BtSLC25A4 crystal structure) | 1 | 2.83 |
| | | Total Simulation Time | 31.79 μs | |

## Notes

### Competing Interest Statement

The authors have declared no competing interest.

### Summary of Updates

Corrected typos and combined supporting information with main text

## References

1. T. S. Luongo, et al., SLC25A51 is a mammalian mitochondrial NAD+ transporter. Nature 588, 174–179 (2020).

2. N. Kory, et al., MCART1/SLC25A51 is required for mitochondrial NAD transport. Sci. Adv. 6 (2020).

3. E. Girardi, et al., Epistasis-driven identification of SLC25A51 as a regulator of human mitochondrial NAD import. Nat. Commun. 2020 111 11, 1–9 (2020).

4. M. Ziegler, et al., Welcome to the Family: Identification of the NAD+ Transporter of Animal Mitochondria as Member of the Solute Carrier Family SLC25. Biomol. 2021, Vol. 11, Page 880 11, 880 (2021).

5. E. Pebay-Peyroula, et al., Structure of mitochondrial ADP/ATP carrier in complex with carboxyatractyloside. Nat. 2003 4266962 426, 39–44 (2003).

6. J. J. Ruprecht, E. R. S. Kunji, Structural Mechanism of Transport of Mitochondrial Carriers. Annu. Rev. Biochem. 90, 535–558 (2021).

7. A. J. Robinson, C. Overy, E. R. S. Kunji, The mechanism of transport by mitochondrial carriers based on analysis of symmetry. Proc. Natl. Acad. Sci. U. S. A. 105, 17766–17771 (2008).

8. J. J. Ruprecht, et al., Structures of yeast mitochondrial ADP/ATP carriers support a domain-based alternating-access transport mechanism. Proc. Natl. Acad. Sci. U. S. A. 111, E426–E434 (2014).

9. H. Nury, et al., Structural basis for lipid-mediated interactions between mitochondrial ADP/ATP carrier monomers. FEBS Lett. 579, 6031–6036 (2005).

10. R. Springett, M. S. King, P. G. Crichton, E. R. S. Kunji, Modelling the free energy profile of the mitochondrial ADP/ATP carrier. Biochim. Biophys. Acta - Bioenerg. 1858, 906–914 (2017).

11. M. S. King, M. Kerr, P. G. Crichton, R. Springett, E. R. S. Kunji, Formation of a cytoplasmic salt bridge network in the matrix state is a fundamental step in the transport mechanism of the mitochondrial ADP/ATP carrier. 1857, 14–22 (2016).

12. S. Todisco, G. Agrimi, A. Castegna, F. Palmieri, Identification of the Mitochondrial NAD+ Transporter in Saccharomyces cerevisiae*. J. Biol. Chem. 281, 1524–1531 (2006).

13. F. Palmieri, et al., Molecular identification and functional characterization of Arabidopsis thaliana mitochondrial and chloroplastic NAD+ carrier proteins. J. Biol. Chem. 284, 31249–31259 (2009).

14. F. Palmieri, The mitochondrial transporter family SLC25: Identification, properties and physiopathology. Mol. Aspects Med. 34, 465–484 (2013).

15. X. A. Cambronne, et al., Biosensor reveals multiple sources for mitochondrial NAD+. Science (80-.). 352, 1474–1477 (2016).

16. J. Jumper, et al., Highly accurate protein structure prediction with AlphaFold. Nat. 2021 5967873 596, 583–589 (2021).

17. A. Waterhouse, et al., SWISS-MODEL: Homology modelling of protein structures and complexes. Nucleic Acids Res. 46, W296–W303 (2018).

18. C. J. Williams, et al., MolProbity: More and better reference data for improved all-atom structure validation. Protein Sci. 27, 293–315 (2018).

19. G. Studer, et al., QMEANDisCo-distance constraints applied on model quality estimation. Bioinformatics 36, 1765–1771 (2020).

20. E. M. Mejia, G. M. Hatch, Mitochondrial phospholipids: role in mitochondrial function. J. Bioenerg. Biomembr. 2015 482 48, 99–112 (2015).

21. J. Comte, B. Maǐsterrena, D. C. Gautheron, Lipid composition and protein profiles of outer and inner membranes from pig heart mitochondria. Comparison with microsomes. BBA - Biomembr. 419, 271–284 (1976).

22. S. Jo, T. Kim, V. G. Iyer, W. Im, CHARMM-GUI: A web-based graphical user interface for CHARMM. J. Comput. Chem. 29, 1859–1865 (2008).

23. J. Lee, et al., CHARMM-GUI Input Generator for NAMD, GROMACS, AMBER, OpenMM, and CHARMM/OpenMM Simulations Using the CHARMM36 Additive Force Field. J. Chem. Theory Comput. 12, 405–413 (2016).

24. E. L. Wu, et al., CHARMM-GUI membrane builder toward realistic biological membrane simulations. J. Comput. Chem. 35, 1997–2004 (2014).

25. J. J. Ruprecht, et al., The Molecular Mechanism of Transport by the Mitochondrial ADP/ATP Carrier. Cell 176, 435–447.e15 (2019).

26. A. L. Duncan, J. J. Ruprecht, E. R. S. Kunji, A. J. Robinson, Cardiolipin dynamics and binding to conserved residues in the mitochondrial ADP/ATP carrier. Biochim. Biophys. Acta - Biomembr. 1860, 1035–1045 (2018).

27. P. G. Crichton, et al., Trends in thermostability provide information on the nature of substrate, inhibitor, and lipid interactions with mitochondrial carriers. J. Biol. Chem. 290, 8206–8217 (2015).

28. N. Senoo, et al., Cardiolipin, conformation, and respiratory complex-dependent oligomerization of the major mitochondrial ADP/ATP carrier in yeast. Sci. Adv. 6, 780–808 (2020).

29. J. M. Eller, et al., Flow Cytometry Analysis of Free Intracellular NAD+ Using a Targeted Biosensor. Curr. Protoc. Cytom. 88, e54 (2019).

30. D. R. Nelson, C. M. Felix, J. M. Swanson, Highly conserved charge-pair networks in the mitochondrial carrier family. J. Mol. Biol. 277, 285–308 (1998).

31. D. V. Miniero, M. Monné, M. A. Di Noia, L. Palmieri, F. Palmieri, Evidence for Non-Essential Salt Bridges in the M-Gates of Mitochondrial Carrier Proteins. Int. J. Mol. Sci. 2022, Vol. 23, Page 5060 23, 5060 (2022).

32. Q. Yi, et al., Molecular dynamics simulations on apo ADP/ATP carrier shed new lights on the featured motif of the mitochondrial carriers. Mitochondrion 47, 94–102 (2019).

33. C. L. Pierri, F. Palmieri, A. De Grassi, Single-nucleotide evolution quantifies the importance of each site along the structure of mitochondrial carriers. Cell. Mol. Life Sci. 71, 349–364 (2014).

34. S. Forli, et al., Computational protein–ligand docking and virtual drug screening with the AutoDock suite. Nat. Protoc. 2016 115 11, 905–919 (2016).

35. G. M. Morris, et al., Software news and updates AutoDock4 and AutoDockTools4: Automated docking with selective receptor flexibility. J. Comput. Chem. 30, 2785–2791 (2009).

36. E. R. S. Kunji, A. J. Robinson, The conserved substrate binding site of mitochondrial carriers. Biochim. Biophys. Acta - Bioenerg. 1757, 1237–1248 (2006).

37. A. J. Robinson, E. R. S. Kunji, Mitochondrial carriers in the cytoplasmic state have a common substrate binding site. Proc. Natl. Acad. Sci. U. S. A. 103, 2617–2622 (2006).

38. E. R. S. Kunji, M. Harding, Projection structure of the atractyloside-inhibited mitochondrial ADP/ATP carrier of Saccharomyces cerevisiae. J. Biol. Chem. 278, 36985–36988 (2003).

39. Y. Wang, E. Tajkhorshid, Electrostatic funneling of substrate in mitochondrial inner membrane carriers. Proc. Natl. Acad. Sci. U. S. A. 105, 9598–9603 (2008).

40. D. Heidkämper, V. Müller, D. R. Nelson, M. Klingenberg, Probing the role of positive residues in the ADP/ATP carrier from yeast. The effect of six arginine mutations on transport and the four ATP versus ADP exchange modes. Biochemistry 35, 16144–16152 (1996).

41. M. Monné, F. Palmieri, E. R. S. Kunji, The substrate specificity of mitochondrial carriers: Mutagenesis revisited. http://dx.doi.org/10.3109/09687688.2012.737936 30, 149–159 (2013).

42. E. M. Krammer, et al., High-Chloride Concentrations Abolish the Binding of Adenine Nucleotides in the Mitochondrial ADP/ATP Carrier Family. Biophys. J. 97, L25–L27 (2009).

43. F. Dehez, E. Pebay-Peyroula, C. Chipot, Binding of ADP in the mitochondrial ADP/ATP carrier is driven by an electrostatic funnel. J. Am. Chem. Soc. 130, 12725–12733 (2008).

44. V. Mavridou, et al., Substrate binding in the mitochondrial ADP/ATP carrier is a step-wise process guiding the structural changes in the transport cycle. Nat. Commun. 2022 131 13, 1–12 (2022).

45. H. Majd, et al., Screening of candidate substrates and coupling ions of transporters by thermostability shift assays. Elife 7 (2018).

46. J. Mifsud, et al., The substrate specificity of the human ADP/ATP carrier AAC1. http://dx.doi.org/10.3109/09687688.2012.745175 30, 160–168 (2013).

47. D. H. Williamson, P. Lund, H. A. Krebs, “The Redox State of Free Nicotinamide-Adenine Dinucleotide in the Cytoplasm and Mitochondria of Rat Liver” (1967).

48. Q. Zhang, D. W. Piston, R. H. Goodman, Regulation of corepressor function by nuclear NADH. Science (80-.). 295, 1895–1897 (2002).

49. Y. P. Hung, J. G. Albeck, M. Tantama, G. Yellen, Imaging cytosolic NADH-NAD + redox state with a genetically encoded fluorescent biosensor. Cell Metab. 14, 545–554 (2011).

50. Y. Zhao, et al., Genetically encoded fluorescent sensors for intracellular NADH detection. Cell Metab. 14, 555–566 (2011).

51. M. Schlame, Protein crowding in the inner mitochondrial membrane. Biochim. Biophys. Acta - Bioenerg. 1862, 148305 (2021).

52. Y. Lee, C. Willers, E. R. S. Kunji, P. G. Crichton, Uncoupling protein 1 binds one nucleotide per monomer and is stabilized by tightly bound cardiolipin. Proc. Natl. Acad. Sci. U. S. A. 112, 6973–6978 (2015).

53. S. Ghosh, et al., An essential role for cardiolipin in the stability and function of the mitochondrial calcium uniporter. Proc. Natl. Acad. Sci. U. S. A. 117, 16383–16390 (2020).

## SI References

1. J. S. Papadopoulos, R. Agarwala, COBALT: constraint-based alignment tool for multiple protein sequences. Bioinformatics 23, 1073–1079 (2007).

2. K. Okonechnikov, et al., Unipro UGENE: A unified bioinformatics toolkit. Bioinformatics 28, 1166–1167 (2012).

3. A. Waterhouse, et al., SWISS-MODEL: Homology modelling of protein structures and complexes. Nucleic Acids Res. 46, W296–W303 (2018).

4. E. Pebay-Peyroula, et al., Structure of mitochondrial ADP/ATP carrier in complex with carboxyatractyloside. Nat. 2003 4266962 426, 39–44 (2003).

5. J. J. Ruprecht, et al., The Molecular Mechanism of Transport by the Mitochondrial ADP/ATP Carrier. Cell 176, 435–447.e15 (2019).

6. J. Jumper, et al., Highly accurate protein structure prediction with AlphaFold. Nat. 2021 5967873 596, 583–589 (2021).

7. C. J. Williams, et al., MolProbity: More and better reference data for improved all-atom structure validation. Protein Sci. 27, 293–315 (2018).

8. G. Studer, et al., QMEANDisCo-distance constraints applied on model quality estimation. Bioinformatics 36, 1765–1771 (2020).

9. S. Jo, T. Kim, W. Im, Automated Builder and Database of Protein/Membrane Complexes for Molecular Dynamics Simulations. PLoS One 2, e880 (2007).

10. E. L. Wu, et al., CHARMM-GUI membrane builder toward realistic biological membrane simulations. J. Comput. Chem. 35, 1997–2004 (2014).

11. J. Comte, B. Maǐsterrena, D. C. Gautheron, Lipid composition and protein profiles of outer and inner membranes from pig heart mitochondria. Comparison with microsomes. BBA - Biomembr. 419, 271–284 (1976).

12. M. Sathappa, N. N. Alder, The ionization properties of cardiolipin and its variants in model bilayers. Biochim. Biophys. Acta - Biomembr. 1858, 1362–1372 (2016).

13. G. Olofsson, E. Sparr, Ionization Constants pKa of Cardiolipin. PLoS One 8 (2013).

14. Bekker H., et al., Gromacs: A parallel computer for molecular dynamics simulations. Physics computing 92. World Scientific, Singapore, pp. 252–256.

15. M. J. Abraham, et al., GROMACS: High performance molecular simulations through multi-level parallelism from laptops to supercomputers. SoftwareX 1–2, 19–25 (2015).

16. T. S. Luongo, et al., SLC25A51 is a mammalian mitochondrial NAD+ transporter. Nature 588, 174–179 (2020).

17. J. M. Eller, et al., Flow Cytometry Analysis of Free Intracellular NAD+ Using a Targeted Biosensor. Curr. Protoc. Cytom. 88, e54 (2019).

