## Supplementary figures and images for "Dynamics of Mitochondrial NAD^+^ Import Reveal Preference for Oxidized Ligand and Substrate Led Transport"

### Movie2

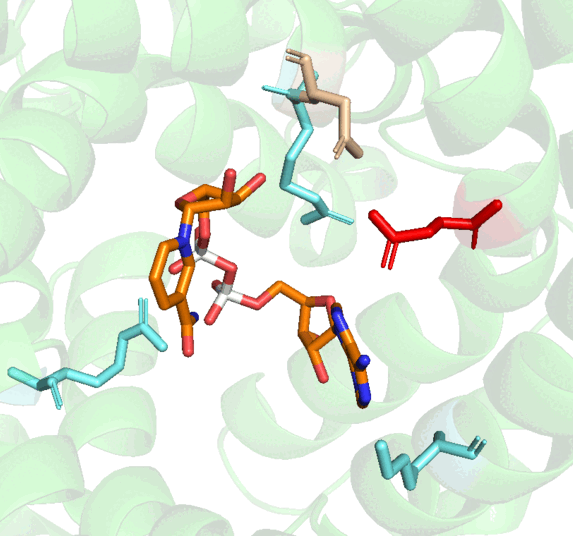
